# Nerve injury promotes glial immune responses through a Draper/Ninjurin A pathway

**DOI:** 10.1101/2025.07.21.665563

**Authors:** Cole R. Brashaw, Annie M. Griffin, Sean D. Speese, Mary A. Logan

## Abstract

Degenerating neurons elicit striking immune reactions from glial cells, including directed invasion of injury sites and engulfment of neuronal debris. While these conserved glial immune responses are neuroprotective, our mechanistic understanding of glial immunity in the damaged and diseased brain is still incomplete. Here, using an *in vivo* nerve injury assay in the adult *Drosophila* olfactory system, we characterize a novel role for the transmembrane adhesion molecule Ninjurin A (NijA). We show that NijA is transcriptionally upregulated in neuropil ensheathing glia, but not local astrocytes, within hours after olfactory nerve transection. In *NijA* mutants, glia fail to properly infiltrate areas that contain severed olfactory nerves, and degenerating axonal debris is not cleared from the CNS. One well-defined signaling cascade critical for ensheathing glial clearance of damaged olfactory axons is the conserved MEGF10/Draper pathway, which includes the engulfment receptor Draper, downstream transcriptional complex AP-1, and known gene target MMP-1. We show that injury-induced transcription of *NijA* in responding glia requires the Draper receptor but is independent of MMP-1, suggesting a parallel signaling cascade is activated downstream of Draper in responding glia. Our findings reveal an essential role for the glial adhesion factor NijA in morphological and phagocytic responses to CNS damage, highlighting this conserved molecule as a new potential glial therapeutic target for neurodegenerative conditions.

## Introduction

As the primary immune surveyors of the brain, glial cells respond quickly to neural trauma while altering gene expression, migrating to areas of damage, and phagocytosing degenerating cells and projections (Barres, 2008; Magaki et al., 2018; Raiders et al., 2021). These responses offer neuroprotection by sequestering and removing neurotoxic cell fragments while continuing to support intact adjacent neurons (Logan and Freeman, 2007; Allen and Barnes, 2009; Jäkel and Dimou, 2017; Jung and Chung, 2018; Yu, et al., 2022). Notably, dysfunctional glial immunity likely bolsters neurodegenerative disease states and related cognitive decline (Sokolowski and Mandell 2011; Mosher et al., 2014; Purice et al., 2016; Bussian et al., 2018; Donnelly et al., 2020). A comprehensive understanding of the signaling pathways and communication relays of glial immune behaviors is essential to identify candidate therapeutic intervention points in the context of CNS trauma and disease.

Glial immune responses to injury are remarkably conserved at cellular and molecular levels, including defined transcriptional programs, activated kinase cascades, and phagocytic engulfment programs. *Drosophila* provides a powerful *in vivo* system to investigate innate glial responses to neurodegeneration (Lye and Chtarbanova, 2018; Kim, et al., 2020). Severed nerves undergo a classic Wallerian degeneration program in adult flies (Freeman, 2006), triggering conserved immune pathways in local glial cells (Lee and Sun, 2015; Logan, 2017; Hilu-Dadia and Kurant, 2020). This includes activation of Draper/MEGF10, a well-defined engulfment receptor that initiates critical signaling cascades in responding glia (Logan et al., 2012; MacDonald et al., 2013; Doherty et al., 2014; Purice et al., 2017; Wu et al., 2009; Scheib et al., 2012; Chung et al., 2013). Activated Draper initiates c-Jun N-terminal kinase (JNK) pathway signaling and the JNK activator protein-1 (AP-1) transcriptional complex, a heterodimer of Jra and Kayak, homologous to mammalian c-Jun and c-Fos respectively (Riesgo-Escovar and Hafen, 1997; Parks et al., 2004; Pastor-Pareja et al., 2004, Page-McCaw et al., 2007; MacDonald et al., 2013; Doherty et al., 2014). AP-1 activity also converges with signal transducer and activator of transcription (STAT92E) to upregulate the transcriptional target matrix metalloproteinase-1 (*MMP-1*) in glia responding to degenerating axons (Logan et al., 2012; MacDonald et al., 2013; Doherty et al., 2014; Purice et al., 2017). MMPs come in secreted and membrane-tethered forms. Due to their endopeptidase activity, they are heavily implicated in cell migration and remodeling, in part due to their ability to cleave extracellular matrix (ECM), transmembrane receptors, and other relevant substrates (Nagase and Woessner, 1999; Vu and Werb, 2000; Visse and Nagase, 2003; Page-McCaw, 2007; Kobayashi et al., 2008; Zitka et al., 2010; LaFever et al., 2017; Chen and Parks, 2019; Pan et al., 2022). However, little is known about MMP function and specific cleavage targets that facilitate glial motility and phagocytic clearance of damaged neurons in the injured brain.

In a previous RNA-seq screen, we identified *Ninjurin A (NijA*) as a candidate upregulated gene in the adult *Drosophila* ventral nerve cord following transection of leg and wing nerve projections (Purice et al., 2017). Vertebrate Ninjurin-1 (Nerve injury-induced protein 1) was first identified in a screen for upregulated factors in rodent sciatic nerve injury following transection or crush injury (Araki and Milbrandt, 1996). Since then, Ninjurin induction has been observed in other models of injury and inflammation (Lee et al., 2016; Weerasinghe-Mudiyanselage et al., 2021; Jennewein et al., 2015; Zhu and Xu, 2025), as well as some cell migration/outgrowth developmental processes (Zhang et al., 2006, Kwon et al., 2024). Notably, targeted MMP cleavage of Ninjurin proteins has been described in flies and mammals. For example, macrophage-expressed NINJ-1 can be cleaved by MMP-9 to release the secreted ectodomain of NINJ-1, which attenuates inflammation (Jeon et al., 2020). In flies, MMP-1 cleaves NijA to promote extension of growing tracheal networks (Zhang et al., 2006) and migration of sequestered haemocytes in developing larvae (Kwon et al., 2024). However, it is unclear precisely how Ninjurins contribute to glial responses to acute nerve injury.

Here, we investigated the role of NijA in glia responding to degenerating axons in the adult olfactory system. We report that NijA is acutely upregulated in local ensheathing glia and is required for proper glial infiltration of injured CNS tissue and for timely phagocytic clearance of severed axons. We also show that injury-induced upregulation of NijA requires the glial immune receptor Draper and propose that NijA is a novel component of the Draper signaling pathway activated in parallel to the AP-1/MMP-1 transcriptional cascade in reactive glia.

## Results

### 2.1. Ninjurin A is upregulated in the central Drosophila brain after antennal nerve transection

Through previous screening efforts, we demonstrated that *NijA* transcription is significantly elevated in ventral nerve cord tissue following peripheral nerve injury; however, the specific cell types expressing *NijA* after axotomy remained unknown (Purice et al., 2017). Turning to the more accessible olfactory system, we asked if *NijA* is upregulated in glia in response to degenerating axons. Surgical ablation of the external antennae and or/maxillary palp (MP) structures severs nerves that contain olfactory receptor neuron (ORN) axons projecting to the antennal lobes (ALs) of the central brain (Bhandawat et al., 2010; Mosca and Luo, 2014) (Figure 1A & B). In uninjured brains, pan-glial expression of membrane-tethered GFP, combined with immunostaining for the pre-synaptic molecule bruchpilot (brp), shows glial membranes surrounding the ALs and neuronal cell bodies in adjacent cortex (Figure 1C). We performed bilateral ablation of the antennae, followed by Hybridization Chain Reaction (HCR) (Choi et al., 2014; Choi et al., 2018) to visualize *NijA* transcripts and compared uninjured, 2 hours post injury (hpi), 4 hpi, 8 hpi, and 16hpi (Figure 1D-H, Figure S1A). In uninjured brains, *NijA* was undetectable in the ALs and the surrounding cortex (Figure 1D). At 2hpi, *NijA* transcripts were observed at the periphery of the ALs and, to a lesser extent, in the cortex (Figure 1E). Larger, higher intensity puncta at the AL edges likely represent newly forming transcripts accumulating within nuclei or cell bodies (arrows, Figure 1E & E’). NijA levels increased further at 4 hpi and 8 hpi broadly around the ALs and in adjacent cortex (Figure 1F & G) with *NijA* transcripts consistently visible in the interior of the ALs at 8hpi (Figure 1G, brackets), suggesting they are transported to distal glial processes. Finally, *NijA* levels were notably reduced by 16hpi, as compared to 4 hpi and 8 hpi (Figure 1H). We observed a similar pattern of induction in NijA-GFP gene trap animals (BL59604). Little to no GFP was present in the brains of uninjured animals, but increased GFP was detected around ALs and in adjacent cortex 3 days after nerve injury (Figure 1I). In addition, no *NijA* HCR signal was detected in the central brains of *NijA* null animals at 4hpi or 16hpi (Figure 1K). RT-qPCR confirmed lack of *NijA* expression in the null mutant strain (Figure S1B).

**Figure 1.**
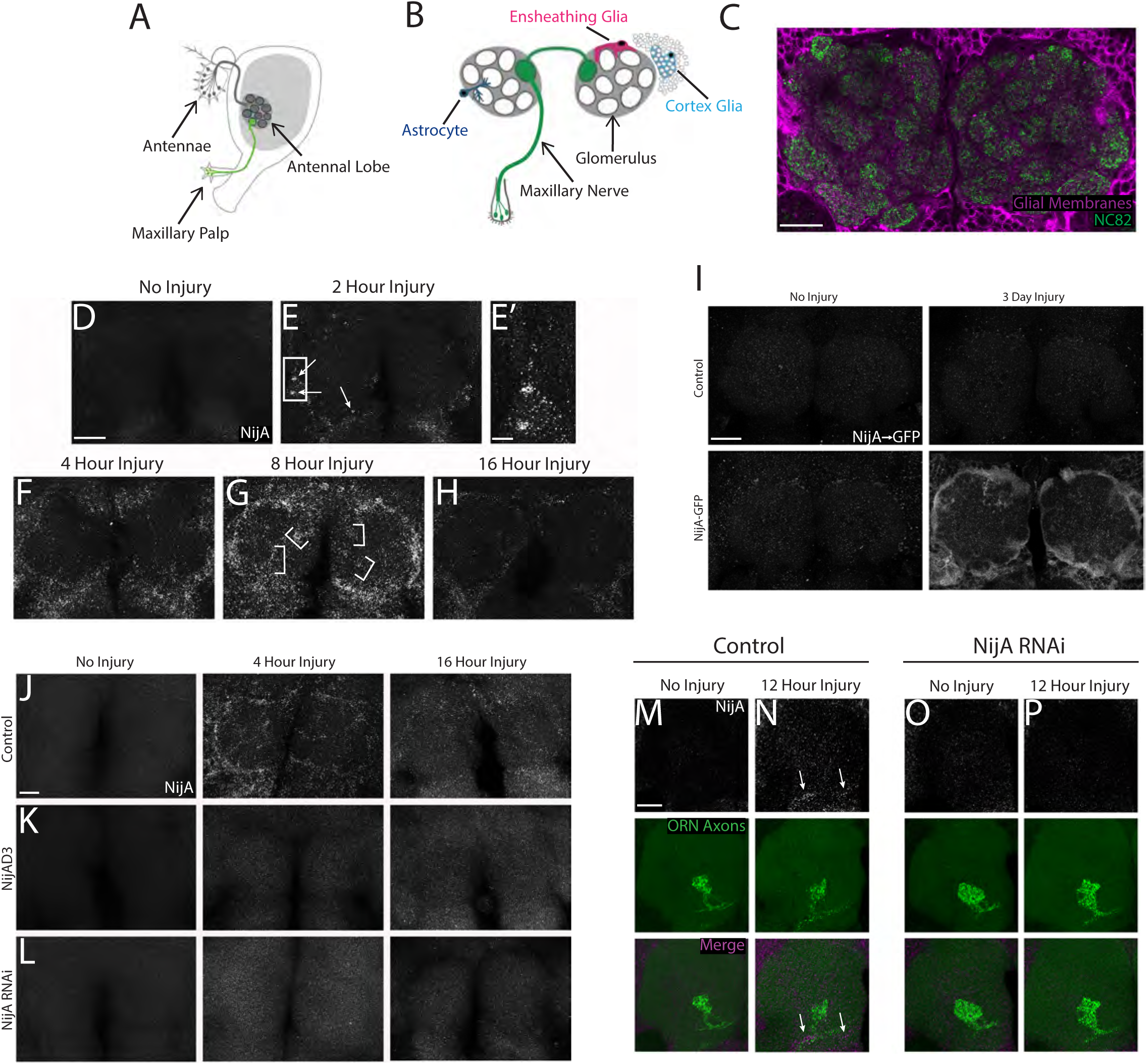
NijA is upregulated in the central brain after olfactory nerve axotomy. **(A)** Side view *Drosophila* head schematic shows olfactory sensory organs, antennae and maxillary palps (MPs), which contain olfactory receptor neurons (ORNs) that project to the antennal lobes (ALs) of the central brain. MP ORNs are labeled green. **(B)** Cartoon frontal view of antennal lobe region in the central brain. MP ORNs (green) project through MP nerves to innervate discrete glomeruli. The AL neuropil contains ensheathing glia (EG) (magenta) and astrocytes (dark blue). Neuronal cell bodies in the cortex (black circles) are enwrapped by cortex glia (light blue). **(C)** Single (1µm) confocal slice of antennal lobe region. Glial membranes are genetically labelled with GFP (pseudocolored magenta). AL glomeruli labeled with pre-synaptic marker NC82 (anti-brp, green). Genotype: *w^1118^; UAS-mCD8::GFP/+; repo-gal4/+*. **(D-H)** Confocal Z-stack maximum intensity projections (10µm) of *NijA* transcripts labeled by hybridization chain reaction (HCR) in AL regions at 0 hpi **(D)**, 2 hpi **(E)**, 4 hpi **(F)**, 8 hpi **(G)** and 16 hpi **(H)**. Arrows in **(E)** point to large puncta consistent with newly forming NijA transcripts accumulating in nuclei and/or cytoplasm at 2hpi. **(E’)** High magnification image of boxed region shown in **(E)**. Brackets in **(G)** denote representative regions of increased *NijA* signal in the AL interior. Genotype: *w^1118^*. **(I)** Confocal Z-stack maximum intensity projections (5µm) of control and *NijA* GFP gene trap flies at 0 and 3 dpi. Brains stained for anti-GFP (green). Genotypes: control= *w^1118^*. NijA-GFP = *y^1^w\**; *Mi{MIC}NijA^MI15170^*. **(J-L)** Representative images of *NijA* HCR in control, *NijA* null (NijAD3), and pan-glial *NijA* knockdown (NijA RNAi) brains at 0 hpi, 4 hpi, and 16 hpi. Confocal Z-stack maximum intensity projections (10µm) shown. Genotypes: control = *w^1118^; repo-gal4/+.* NijAD3 = *w^1118^; NijAD3/NijAD3*. NijA RNAi = *w^1118^; NijA^RNAi50532^/+; repo-gal4/+*. **(M-P)** Confocal Z-stack maximum intensity projections (5µm) of *NijA* HCR (grayscale) and GFP-labeled OR85e axons (green) at 0 hpi **(M, O)** and 12 hpi **(N, P)**. Arrows in **(N)** point to *NijA* HCR signal localized to GFP^+^ maxillary nerve axons. Genotypes: control = *w^1118^; OR85e-mCD8::GFP/+; repo-gal4/+.* NijA RNAi = *w^1118^; OR85e-mCD8::GFP/NijA^RNAi50632^; repo-Gal4/+*. Scale bars for all panels except E’= 30µm. Scale bar in E’ = 8µm. hpi = hours post-injury. dpi = days post-injury.

While antennal ablation severs most axons projecting to the ALs, the maxillary palps (MPs) house only ∼15% of adult ORNs, which extend to 5 out of ∼50 glomeruli in each AL (de Bruyne et al., 1999; Verschut et al., 2018; Horne et al., 2018). To determine if *NijA* is induced locally in response to smaller, focal injury, we performed bilateral MP ablation and *NijA* HCR on flies expressing membrane-GFP in a subset of MP ORNs (*OR85e-mCD8::GFP*). At 12 hpi, we observed notable *NijA* signal along injured MP nerves that was not present in uninjured animals (Figure 1M & N). No increase in *NijA* HCR signal was detected around the ALs or the surrounding cortex (Figure 1N), as we consistently saw after severing the larger antennal nerves (Figure 1E & H), suggesting that *NijA* upregulation is induced locally by degenerating axons and proportionate to the size of the injury.

### 2.2. Glial knockdown of NijA leads to delayed clearance of adult degenerating axons and attenuated actin dynamics after injury

*NijA* upregulation after axotomy appeared consistent with a glial expression pattern. We first used the pan-glial driver repo-Gal4 to knockdown *NijA* by RNAi and performed confirmational comparative RT-qPCR analysis for *UAS-NijA^RNAi^* transgenic lines (Figure S1C). NijA was significantly reduced in flies carrying the *UAS-NijA^RNAi50632^*transgene but unchanged by *UAS-NijA^RNAi51358^* (Figure S1C). Thus, BL50632 was used for all subsequent *in vivo* RNAi experiments. Next, we performed antennal ablations, followed by *NijA* HCR at 4 hpi and 16 hpi in glial *NijA* knockdown animals. As with *NijA* null mutants, we observed no *NijA* upregulation at either time point after axotomy (Figure 1L), indicating that the acute upregulation of *NijA* after transection of olfactory nerves projecting into the central brain occurs exclusively in glia in this brain region.

To explore a functional role for *NijA*, we first assessed clearance of degenerating ORN axons. We knocked down glial *NijA* in flies expressing membrane-tethered GFP in MP ORNs (*OR85e-mCD8::GFP*) and quantified GFP+ neuronal debris 1 day after maxillary and antennal nerve axotomy. Significantly more GFP+ debris remained in *NijA* knockdown brains as compared to controls (Figure 2A & B), suggesting that *NijA*-deficient glia do not properly clear degenerating axons.

**Figure 2.**
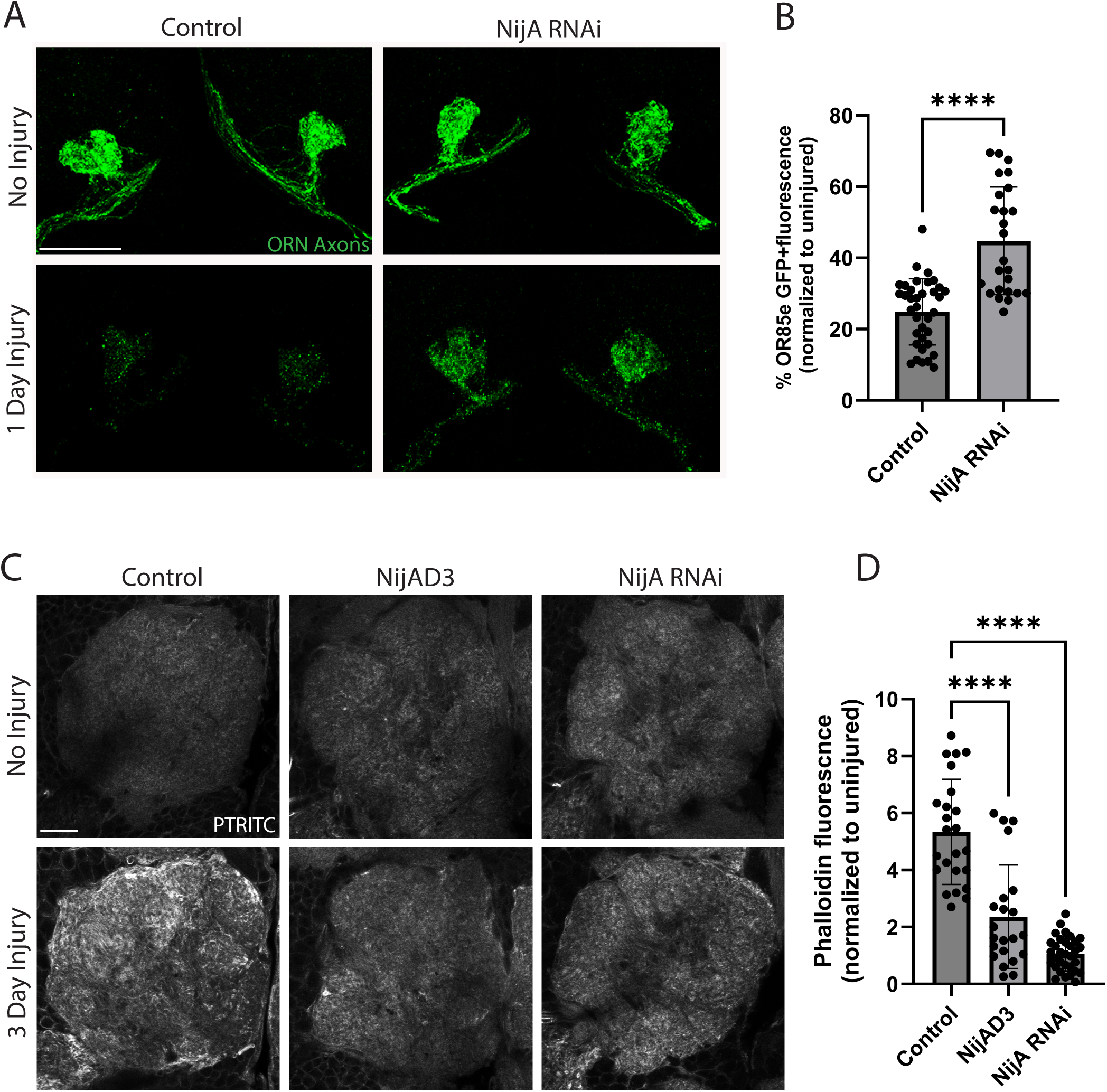
NijA is required for glial recruitment to injury sites and clearance of degenerating ORN axons. **(A)** Representative confocal Z-stack maximum intensity projections (16µm) high magnification images of GFP-labeled maxillary ORN axons (green) in uninjured versus 1 dpi, comparing control and pan-glial *NijA* knockdown (NijA RNAi) animals. Genotypes: control = *w^1118^; OR85e-mCD8::GFP/+; repo-gal4/+*. NijA RNAi = *w^1118^; OR85e-mCD8::GFP/NijA^RNAi50632^; repo-Gal4/+*. **(B)** Quantification of GFP-labeled projections in OR85e glomeruli 1 day after injury (experiment shown in **A**). ALs quantified: Control N= 36, NijA RNAi N= 26; Mean ± SD plotted, ****p<0.0001; unpaired t-test. **(C)** Representative images of AL Phalloidin-TRITC immunostainings (pTRITC) (pseudocolored grayscale) in control, *NijA* null (NijAD3), and pan-glial *NijA* knockdown (NijA RNAi) flies at 0 and 3 dpi. Single 1um slices shown. Genotypes: Control = *w^1118^; repo-Gal4/+.* NijAD3 = *w^1118^; NijAD3/NijAD3*. NijA RNAi = *w^1118^; NijA^RNAi50632^/+; repo-gal4/+*. **(D)** Quantification of total AL pTRITC fluorescence in experiments shown in **(C)**. ALs quantified: Control N= 24, NijAD3 N= 22, NijA RNAi N= 32. Mean ± SD plotted;****p<0.0001; Ordinary one-way ANOVA with Dunnett’s *post-hoc* test. Scale bars= 30µm. dpi = days post-injury.

To clear ORN axonal debris in a timely manner, glia must infiltrate the antennal lobes by extending membrane projections throughout the dense AL neuropil tissue (MacDonald et al., 2006; Logan and Freeman, 2007; Doherty et al., 2009; Tewari et al., 2020; Painter, 2008; Purice et al., 2017). To assess glial membrane recruitment to severed nerves, we stained brains with Phalloidin-TRITC (PTRITC), which labels filamentous actin and serves as a proxy for active cytoskeletal remodeling or migration (Leyssen et al., 2005; Winfree et al., 2017; Purice et al., 2017). As previously reported (Purice et al. 2017), we saw robust Phalloidin-TRITC signal surrounding and within the antennal lobes 3 days after antennal and maxillary nerve injury in controls, but this response was strongly attenuated in *NijA* null mutant brains (Figure 2C & D). Notably, this phenotype was comparable following glial-specific knockdown of *NijA* by RNAi (Figure 2C & D).

### 2.3. Antennal lobe ensheathing glia, not astrocytes, upregulate NijA in response to nerve injury

The adult antennal lobes contain two major glial cell types: ensheathing glia (EG), which enwrap and support neuronal projections, and astrocytes, which regulate local synaptic signaling (Stork et al., 2012; Freeman, 2015; Corty and Coutinho-Budd, 2023). To date, EG carry out all known immune functions following antennal nerve axotomy, including phagocytic clearance of axon debris. We used either the EG specific driver *Tifr-Gal4* or astrocyte specific driver *alrm-Gal4* to express membrane-tethered GFP in each subtype and then performed *NijA* HCR on brains 4 hpi. We consistently observed *NijA* signal overlapping with EG membranes (Figure 3A, B, & B’), but not astrocyte membranes (Figure 3E, F, & F’). Further, *NijA* upregulation typically observed 4hpi was strongly attenuated in flies expressing NijA^RNAi^ in EG (Figure 3D & D’), with minimal *NijA* transcript signal that was indiscernible from uninjured controls (Figure 3C).

**Figure 3.**
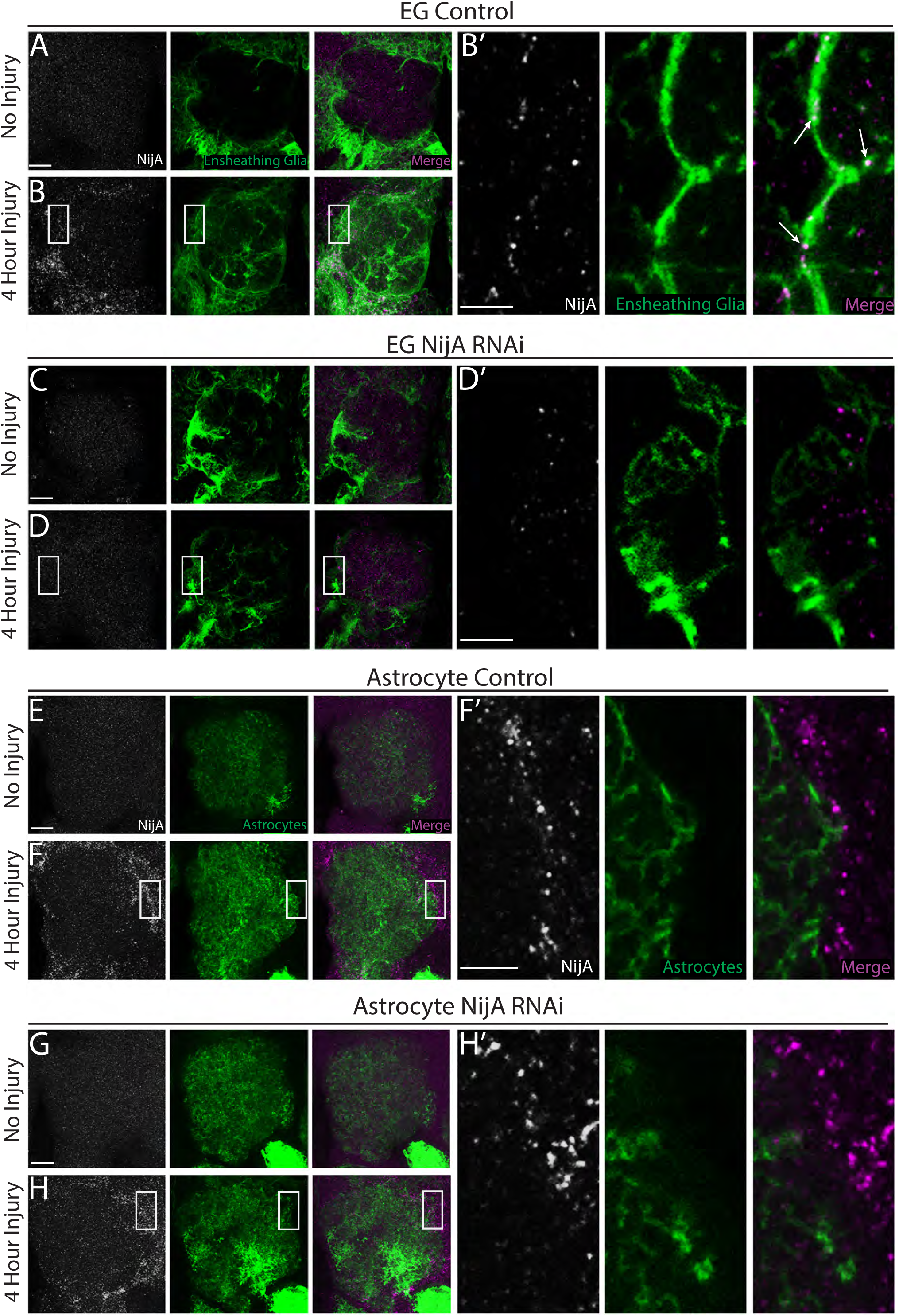
Antennal lobe ensheathing glia, not astrocytes, upregulate NijA in response to olfactory nerve injury. **(A-D)** Representative confocal images of single ALs. EG expressed membrane-tethered GFP. *NijA* transcripts, visualized by HCR, are prevalent at 4 hpi **(B)**. High magnification images of boxed region in **(B)** show characteristic examples of EG membrane and *NijA* overlap (arrows, **B’**). Injury-induced production of *NijA* transcripts is largely absent in flies expressing EG-specific NijA RNAi (C, D, D’). Genotypes: Control = *w^1118^; UAS-mCD8::GFP/+; Tifr-gal4/+.* EG NijA RNAi = *w^1118^; UAS-mCD8::GFP/NijA^RNAi50632^; Tifr-gal4/+*. **(E-H)** Representative confocal images of single ALs. Upregulated *NijA* at 4hpi does not overlap with GFP-labeled astrocyte membranes **(F, F’)**. Astrocyte-specific expression of NijA RNAi does not prevent upregulation or characteristic expression pattern of *NijA* after axotomy **(H, H’)**. Genotypes: Control = *w^1118^; UAS-mCD8::GFP/+; alrm-gal4/+.* Astrocyte NijA RNAi = *w^1118^; UAS-mCD8::GFP/NijA^RNAi50632^; alrm-gal4/+*. Z-stack (10 µm) maximum intensity projections shown in **(A, B, C, D, E, F, G, H)**. High magnification panels **(B’, D’, F’, H’)** are single confocal slice images (1µm). Scale bars = 30 µm **(A, B, C, D, E, F, G, H)** and 5µm **(B’, D’, F’, H’)**.

Adhesion strength of diverse cell types can be altered by either promoting or inhibiting Ninjurin homophilic binding (Araki, 1996; Araki et al., 1997; Ifergan et al., 2011; Kim et al., 2020; Pourmal et al., 2024). It’s well established that EG are exclusively responsible for phagocytosing degenerating ORN axons (Doherty et al., 2009; Su et al., 2013; Musashe et al., 2016) and our findings also point to a role for NijA specifically in EG following antennal nerve injury (Figure 3). However, astrocytic and EG membranes are closely intermingled within antennal lobe glomeruli, and likely even more so as EG extend additional membranes into the AL neuropil after axotomy. Thus, we knocked down *NijA* by RNAi in either EG or astrocytes, severed antennal and maxillary nerves, and stained for Phalloidin-TRITC 3 days post injury. We saw a significant reduction of phalloidin intensity in the ALs post injury in brains expressing NijA^RNAi^ specifically in EG, as compared to controls (Figure 4A & B), while no significant change was quantified following knockdown of *NijA* in astrocytes (Figure 4C & D). This result emphasizes a cell autonomous role for *NijA* in EG and its requirement for proper EG recruitment responses to severed axons. Finally, we knocked down *NijA* in EG cells in *OR85e-mCD8::GFP* flies and quantified significantly more GFP+ axonal debris 1 day after nerve injury as compared to control animals (Figure 4E & F). This supports a requisite role for NijA exclusively in EG injury-activated cytoskeletal events and emphasizes improper recruitment as a factor contributing to delayed debris clearance in NijA-deficient animals.

**Figure 4.**
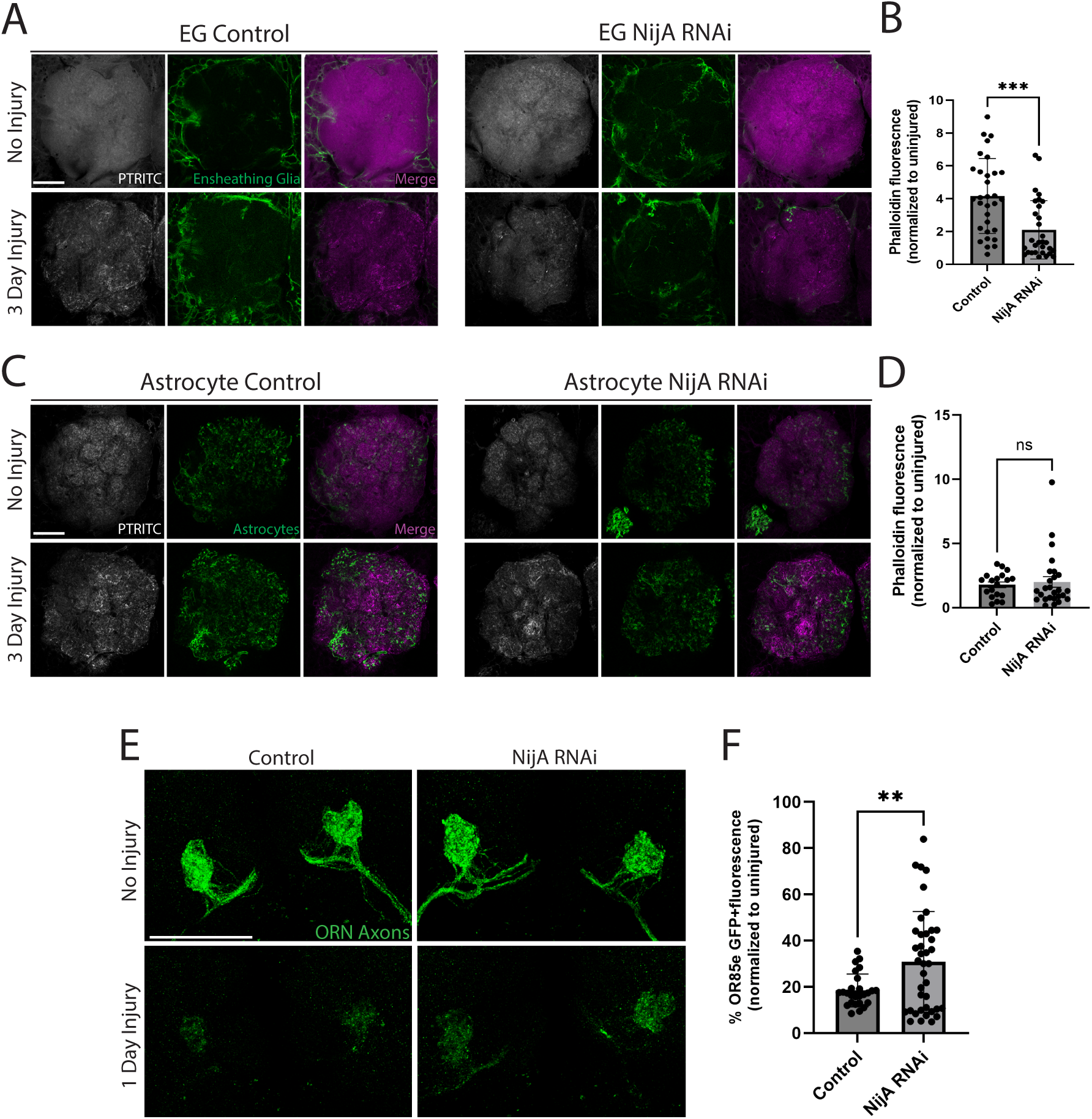
NijA is required in ensheathing glia for injury-induced glial recruitment and clearance of ORN axonal debris. (A) Representative single AL immunostainings of phalloidin (pTRITC) in control and ensheathing glial specific knockdown of *NijA* (EG NijA RNAi) brains, uninjured and 3 dpi. Single 1µm slices shown. Genotypes: Control = *w^1118^; UAS-mCD8::GFP/+; Tifr-gal4/+.* EG NijA RNAi = *w^1118^; UAS-mCD8::GFP/NijA^RNAi50632^; Tifr-gal4/+*. (B) Quantification of total AL pTRITC fluorescence in experiment shown in **(A)**. Control N= 31 ALs, ensheathing glia RNAi N= 30 ALs. Mean ± SD plotted; ***p=0.0002; unpaired t-test. (C) Representative single AL immunostainings of phalloidin (pTRITC) in control and astrocyte specific knockdown of *NijA* (Astrocyte NijA RNAi) brains, uninjured flies and 3 dpi. Single 1um slices shown. Genotypes: Control = *w^1118^; UAS-mCD8::GFP/+; alrm-gal4/+.* Astrocyte NijA RNAi = *w^1118^; UAS-mCD8::GFP/NijA^RNAi50632^; alrm-gal4/+*. (D) Quantification of total AL pTRITC fluorescence in experiment shown in **(C)**. Control N= 19 ALs, astrocyte NijA RNAi N= 26 ALs. Mean ± SD plotted; ns= not significant; unpaired t-test. (E) Confocal images of central AL region with GFP-labeled maxillary ORN axons (green) uninjured and 1 dpi (maxillary palp and antennal nerves) in control animals and ensheathing glial specific *NijA* knockdown flies. Representative Z-stack maximum intensity projections (16µm) shown. Genotypes: Control = *w^1118^; OR85e-mCD8::GFP/+; Tifr-gal4/+.* NijA RNAi = *w^1118^; OR85e-mCD8::GFP/NijA^RNAi50632^; Tifr-gal4/+*. (F) Quantification of GFP^+^ OR85e axon material in glomeruli shown in **(E)**. Control N= 26 glomeruli. NijA RNAi N= 39 glomeruli. Mean ± SD plotted, **p=0.0069; unpaired t-test. Scale bars= 30µm. dpi = days post injury.

### 2.4. Draper is required for NijA upregulation after antennal nerve injury

Ensheathing glial invasion and clearance of degenerating axons after olfactory nerve transection requires the Draper receptor and numerous downstream effectors, including specific transcriptional cascades (Logan et al., 2012; Doherty et al., 2014; Purice et al., 2017). Draper triggers activation of the JNK pathway transcription factor AP-1 which, in turn, boosts draper levels through a positive feedback loop and initiates expression of *MMP1* within hours after axotomy (Doherty et al., 2014; MacDonald et al., 2013; Lu et al., 2017; Purice et al., 2017). Comparing control and draper null mutants, we assessed *NijA* upregulation by HCR following antennal nerve transection at 8 hpi and found that *NijA* induction was blocked in draper mutants (Figure 5A & B). In contrast, glial knockdown of MMP-1 had no discernible effect on injury-induced *NijA* upregulation at 8 hpi (Figure 5C & D).

**Figure 5.**
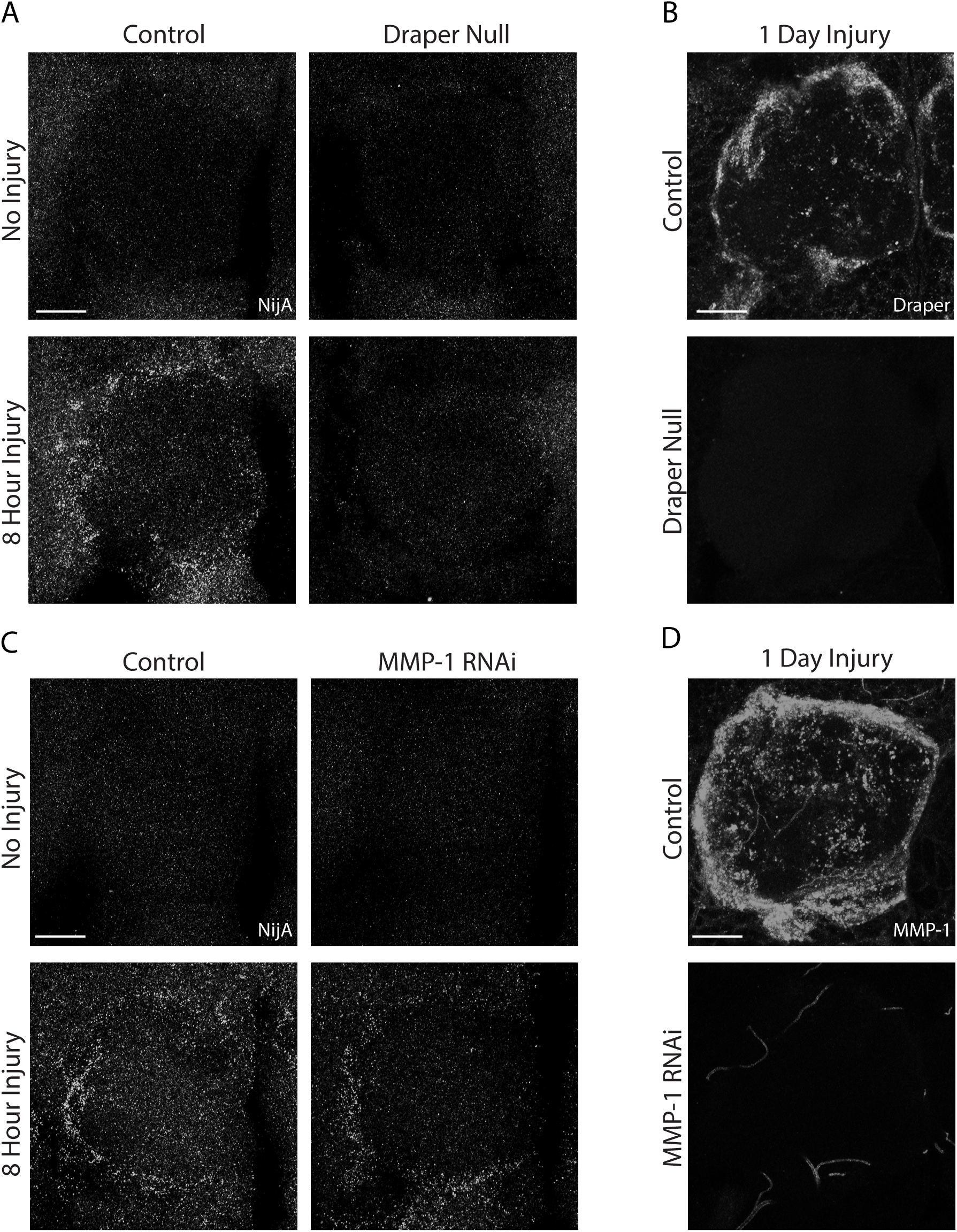
Draper is required for NijA upregulation after olfactory nerve injury. **(A)** Representative Z-stack maximum intensity projections (10µm) of *NijA* transcript in control and Draper null mutants ALs, uninjured and 8 hpi. Single AL shown in each panel. Genotypes: Control = *w^1118^*. Draper null = *w^1118^; draperΔ5rec9*. **(B)** Draper immunostaining in control and *draper* null mutant flies 1 dpi confirms lack of Draper expression. Representative Z-stack maximum intensity projections (10µm) of single AL shown in each panel. Control = *w^1118^*. Draper null = *w^1118^; draperΔ5rec9*. **(C)** Representative Z-stack maximum intensity projections (10µm) of *NijA* transcript in control and glial knockdown of MMP-1 (MMP-1 RNAi) ALs, uninjured and 8 hpi. Single AL shown in each panel. Genotypes: Control = *w^1118^; repo-Gal4/+*. MMP1 RNAi = *w^1118^; repo-Gal4/MMP-1^RNAi^*. **(D)** MMP-1 antibody staining of control and glial MMP-1 RNAi flies at 1dpi confirms lack of MMP-1 in *MMP-1^RNAi^* brains. Representative Z-stack maximum intensity projections (10µm) of single AL shown in each panel. Genotypes: Control = *w^1118^; repo-Gal4/+*. MMP1 RNAi = *w^1118^; repo-Gal4/MMP-1^RNAi^.* Scale bars= 30µm. hpi = hours post injury. dpi = days post injury.

To further investigated NijA function within the Draper signaling pathway, we first asked if NijA is required for injury-induced *draper* upregulation. We transected maxillary nerves and immunostained for Draper at 18 hpi. As previously shown (MacDonald et al., 2006), robust Draper was observed along the maxillary nerve and on maxillary innervated glomeruli in injured control brains (Figure 6A). We detected equivalent upregulation of Draper in injured brains of *NijA* null mutant flies at 18 hpi (Figure 6A & B). Next, we monitored AP-1 activity by using the *in vivo* AP-1 transgenic reporter *Tre-GFP* (Chatterjee and Bohmann, 2012). In control and *NijA* mutant flies, we observed sparse, low levels of GFP in uninjured brains, representing basal AP-1 activity (Chatterjee and Bohmann, 2012; Jemc et al., 2012; Purice et al., 2017), as well as robust upregulation 10 hours after antennal nerve injury (Figure 6C & D). Finally, we assessed injury induction of the AP-1 transcriptional target MMP-1. One day after antennal nerve injury, striking levels of MMP-1 are observed around and throughout the ALs by immunostaining in control brains (Figure 6E) (Purice et al., 2017). Notably, we saw similar robust induction of MMP-1 after injury in flies that lacked *NijA* (Figure 6E & F). Together, these results identify Draper as an upstream receptor required to induce NijA expression in responding glia and suggest that transcriptional regulation of *NijA* falls within an AP-1/MMP-1-independent pathway, although the notion of protein-protein interactions, particularly between MMP-1 and NijA, cannot be excluded.

**Figure 6.**
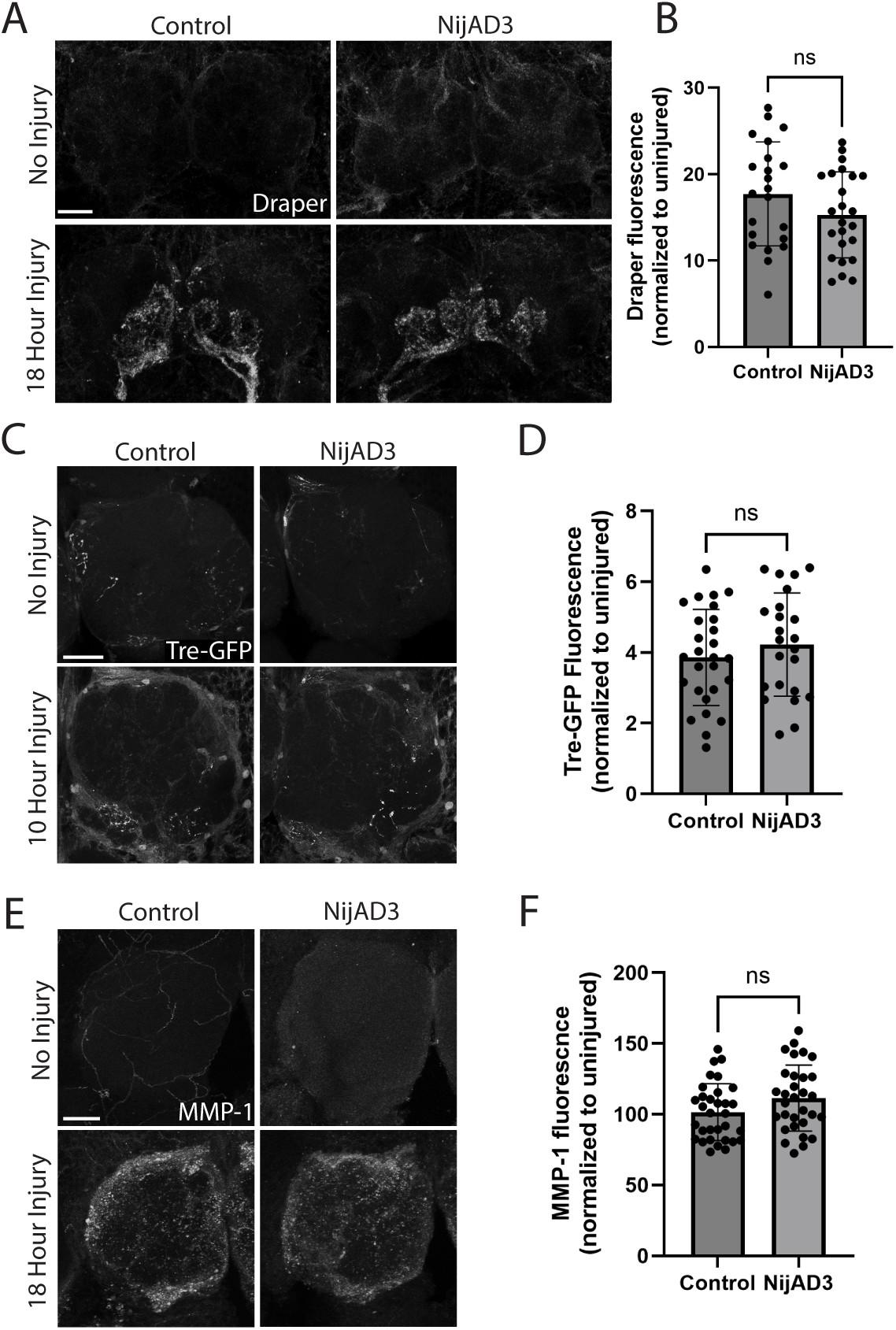
AP-1 activity, Draper, and MMP-1 upregulation are normal in NijA mutant reactive glia. (A) Draper immunostainings in control and *NijA* null mutants (NijAD3), uninjured and 18 hpi (maxillary nerve axotomy). Representative Z-stack maximum intensity projections (20µm) of central AL region containing MP ORN innervated glomeruli shown. Genotypes: Control = *w^1118^*. NijAD3 = *w^1118^; NijAD3/NijAD3*. (B) Quantification of Draper fluorescence in OR85e maxillary palp glomeruli from experiment shown in panel **(A)**. Control N= 22 glomeruli, NijAD3 N= 24 glomeruli. Mean ± SD plotted; ns= not significant. (C) Tre-GFP reporter expression (anti-GFP) of control and *NijA* null mutant (NijAD3) brains, uninjured and 10 hpi (antennal nerve axotomy). Representative Z-stack maximum intensity projections (5µm) of single ALs shown. Genotypes: control = *w^1118^; Tre-GFP/+.* NijAD3 = *w^1118^; Tre-GFP/+; NijAD3/NijAD3*. (D) Quantification of GFP intensity in AL regions from experiment shown in panel **(C)**. Control N= 27 ALs, NijAD3 N= 23 ALs, mean ± SD plotted; ns= not significant. (E) MMP-1 immunostaining of control and *NijA* null mutant (NijAD3) brains, uninjured and 18 hpi. Representative maximum intensity projections (10µm) of single ALs. Genotypes: control = *w^1118^*. NijAD3 = *w^1118^*; *NijAD3/NijAD3*. (F) Quantification of MMP-1 fluorescence in ALs from experiment shown in panel **(E)**. Background signal from uninjured flies was subtracted from analysis. Control N= 32 ALs, NijAD3 N= 31 ALs, mean ± SD plotted; ns= not significant. Scale bars= 30µm. hpi = hours post injury. dpi = days post injury.

## Discussion

Glia sense and respond to a wide range of stressors, including acute trauma, stroke, and other chronic disease-related changes in the CNS. Understanding the orchestration of genetic, metabolic, and morphological transformations of reactive glia is key for identifying candidate points for therapeutic intervention. Here, we characterize a new role for NijA as a positive regulator of ensheathing glial responses to axon injury in the adult Drosophila olfactory system. *NijA* is upregulated in glia within hours after axotomy and is essential for proper glial infiltration of the antennal lobe neuropil and phagocytic clearance of degenerating ORN axons. Finally, we demonstrate that the glial immune receptor Draper is required for injury-induced transcriptional activation of *NijA*. Together, this work defines a new role for NijA as a component of the Draper immune signaling pathway and provides a new molecular foothold for understanding how glia migrate in a targeted manner and clear degenerating projections in the adult CNS.

Ninjurins are small transmembrane proteins with few distinctive motifs. Nonetheless, they are implicated in several immune response events across species. Mammalian *Ninjurin* genes were first noted as transcriptional targets in CNS injury and stroke models (Araki, 1996; Ifergan, et al., 2011; Lee, et al., 2016). These studies support roles for Ninjurins as adhesion molecules in cell migration or tissue remodeling. In *Drosophila*, localization and recruitment of haemocytes (macrophage-like cells) to haematopoietic compartments was recently shown to require NijA (Kwon et al., 2024), supporting a conserved role for NijA adhesion and migration events across species. Our findings highlight a specific role for NijA in facilitating EG membrane extension into dense neuropil regions immediately following nerve injury. Notably, alternative roles for Ninjurins in systemic immunity have been described recently, including polymerization of NINJ-1 on macrophages to facilitate membrane lysis, cell rupture, and death (Kayagaki et al., 2021; Degen et al., 2023). How Ninjurin activity drives cell functions ranging from adhesion to death is unknown, but highlights the need to define how Ninjurin expression, localization, and function are regulated in specific biological contexts.

Very little is known about the specific transcriptional cascades regulating Ninjurin expression in any species. We show that activation of glial Draper is required for *NijA* upregulation after nerve injury, as early as several hours after axotomy. We propose that this occurs in parallel to the known Draper/AP-1/MMP-1 signaling cascade, which is activated in ensheathing glia on a similar timescale (Lu et al., 2017; Purice et al., 2017). Notably, a recent study exploring glial engulfment of pathogenic mutant huntingtin (mHTT) aggregates in the adult olfactory system shows that NijA is required for ensheathing glia to properly internalize mHTT, phenocopying *draper* mutants (Davis et al., 2024). Whether NijA activity is specifically contributing to ORN axon engulfment in our system or if delayed engulfment is a result of poor glial recruitment is not yet clear. In our injury paradigm, it is unlikely that astrocytes fail to upregulate *NijA* because they lack the ability to do so. *NijA* is upregulated in *Drosophila* astrocytes follow impact-induced traumatic brain injury (TBI) (Li et al., 2024), suggesting that glial engagement of *NijA* in conditions of stress or trauma are context and/or region dependent. Questions remain about the signaling cascades and transcriptional regulators upstream of NijA and downstream of Draper in adult glia. In the systemic immune system, *NijA* is a transcriptional target of Abrupt (Ab), a transcription factor primarily known for steroid signaling in various developmental contexts (Kwon, et al., 2024). Ab was not upregulated in our original nerve injury RNA-seq screen (Purice et al., 2017), but this does not exclude the possibility that Ab contributes to *NijA* induction in adult glia in response to stress or trauma.

Although we have focused on NijA and glial immune responses in the antennal lobe (AL) region, which houses all degenerating projections in our injury model, we also observed *NijA* upregulated in adjacent cortex regions after olfactory nerve injury (Figure 1F, G, I). Notably, this cortex signal is lost in *draper* null mutants (Figure 5A). The ALs are compartmentalized, enwrapped by ensheathing glia, and cortex glial cell membranes are never observed in the ALs, even after axotomy. However, Draper is basally expressed in both ensheathing and cortex glia, and, after axotomy, Draper-dependent AP-1 activity is triggered in cortex glia cells adjacent to the ALs (MacDonald et al., 2016; Doherty et al., 2009; Lu et al 2017). This leads to interesting questions about why injury-induced genes are upregulated distally from sites of injury, and if cortex glial specific knockdown could be sufficient to delay axon phagocytosis or ensheathing glial migration or if these represent a broad generalized CNS stress responses in cortex glia following nerve transection.

MMPs are heavily implicated in cell migration and remodeling, in part due to their ability to cleave extracellular matrix (ECM), transmembrane receptors, and other relevant substrates (Chen and Marks, 2009; LaFever et al., 2017), making Ninjurins an intriguing MMP target. Indeed, several lines of evidence support MMP cleavage of NijA in flies. NijA and MMP-1 co-localize at the tracheal cell surface and co-immunoprecipitated at later pre-pupal stages (Zhang et al., 2006). Further, overexpression of MMP-1 in haemocytes replicates the NijA loss-of-function phenotype, while blocking MMP-1 expression mimics *NijA* overexpression (Kwon et al., 2024). Here, we propose that concomitant production of MMP-1 and NijA in ensheathing glia is an important feature of glial infiltration of the AL and clearance of degenerating axons. We find it plausible that MMP-1 cleavage of the NijA N-terminus is important for proper execution of glial migration and/or engulfment of axon debris. Important objectives moving forward include clarifying a putative function for the released extracellular domain of NijA and defining the transcriptional cascades downstream of Draper that trigger *NijA* expression in response to damaged axons.

## Materials and Methods

### Fly stocks and husbandry

For all experiments, sex matched male and female adult flies (between 5 and 15 days old) were raised on a 12-hour light on/off cycle at 25°C. The following transgenic *Drosophila* stocks were used: *w^1118^;OR85e-mCD8::GFP/CyO* (gift from B. Dickson), *UAS-mCD8::GFP* (Bloomington 5137, RRID:BDSC_5137), *repo-Gal4* (MacDonald et al., 2006), *Tifr-Gal4* (Yao et al., 2007), *alrm-Gal4* (Doherty et al., 2009), *Tre-GFP* (Chatterjee and Bohmann, 2012), *draper^Δ5rec9^*(MacDonald et al., 2006), *UAS-Mmp-1^RNAi^* (Uhlirova and Bohmann, 2006), *w^1118^* (Bloomington 5905, RRID:BDSC_5905), *NijAD3* (Gift from A. McCaw), *UAS-NijA^RNAi50632^* (Bloomington 50632, RRID:BDSC_50632), *UAS-NijA^RNAi51358^* (Bloomington 51358, RRID:BDSC_51358), and NijA-GFP gene trap *y^1^ w*; Mi{MIC}NijA^MI15170^* (Bloomington 59604, RRID:BDSC_59604) (Kwon et al., 2024).

### Olfactory Receptor Neuron (ORN) axotomy assay

ORN axotomy was induced by using forceps to surgically remove both third antennal segments and/or maxillary palps of anesthetized flies, severing the antennal and/or maxillary nerves, respectively (McDonald et al., 2006). Flies were maintained on standard food media at 25°C for indicated times until dissection, fixation, and analysis.

### *Drosophila* brain dissection and immunolabeling

Adult *Drosophila* heads were fixed in PFA-PBSTx0.1 (1xPBS, 0.1% Triton X-100, 4% PFA) at room temperature for 20 min. Samples were then washed 1 × 1 min and 2 × 5 min in PBSTx0.01 (1xPBS, 0.01% Triton X-100) rocking at room temperature. Fixed samples were maintained on ice while brains were dissected at room temperature in PBSTx0.01. Tissue was post-fixed in PFA-PBSTx0.1 for 20 min, washed 1×1 min and 2 × 5 min in PBSTx0.1, and incubated overnight with primary antibodies in PBSTx0.1. The next day, samples were washed 1 × 15 and 3 × 30 min with PBSTx0.1 then incubated with secondary antibodies (diluted in PBSTx0.1) for 2 hr at room temperature. Samples were washed 1 × 15 and 3 × 30 min with PBSTx0.1 and then mounted on slides in VECTASHIELD PLUS antifade mounting medium (Vector Labs, H-1900).

### Antibodies

Primary antibodies were used at the following dilutions: chicken anti-GFP (Thermo Fisher, #A10262, RRID:AB_2534023) at 1:1000; mouse anti-Draper (Developmental Studies Hybridoma Bank, 8A1 RRID:AB_2618106 and 5D14 RRID:AB_2618105) at 1:400; mouse anti-MMP-1 (Developmental Studies Hybridoma Bank, 14A3D2 RRID:AB_579782, 3A6B4 RRID:AB_579780, 3B8D12 RRID:AB_579781, 5H7B11 RRID:AB_579779) at 1:50 used at 1:1:1:1 ratio; Phalloidin-TRITC (Sigma, #P1951 RRID:AB_2315148) at 1:250. All secondary antibodies (Jackson Immunoresearch) were used at 1:400: Alexa Fluor 488 donkey anti-chicken (RRID: AB_2340375); Alexa Fluor 488 donkey anti-mouse IgG (RRID: AB_2340846); Alexa Fluor 488 goat anti-mouse IgG2a (RRID: AB_2338855).

### Confocal microscopy and image analysis

All brains were imaged on a Zeiss LSM 700 confocal microscope with a Zeiss 40X 1.4 NA oil immersion lens using Zen software. Brains for each experiment were mounted under a single #1.5 cover glass in VECTASHIELD PLUS and imaged on the same day with identical laser and detector settings. Volocity 3D Image Analysis Software (Perkin Elmer, RRID:SCR_002668) was used for fluorescence quantification. Quantification of OR85e GFP-labeled glomeruli was performed on 3D volumes segmented to GFP signal. To quantify PTRITC, Draper, GFP (Tre-GFP), and MMP-1 levels in the cortex and antennal lobes of adult brains, total intensity measurements were calculated in regions of interest. Background tracheal MMP-1 staining was subtracted from uninjured and injured conditions (Figure 6C & D) GraphPad Prism (RRID:SCR_002798) was used for statistical analysis including one-way ANOVA with Dunnett’s post hoc test and unpaired t-tests.

### Quantitative Reverse Transcriptase-PCR

Total RNA from heads was collected via Total RNA Purification Micro Kit (Norgen Biotek) and subject to DNAse digestion using Ambion DNA-free kit. Complementary DNA was prepared using qScript cDNA SuperMix kit (Quanta Biosciences). Total RNA was quantified using the Qubit RNA HS assay kit and Qubit Fluorometer, and equal amounts of RNA were added to cDNA synthesis reaction. Quantitative gene expression was carried out on an ABI 7,500 Fast Real-Time PCR machine using Taqman master mix (Applied Biosystems) and the following TaqMan assays: Ninjurin A-Dm01798347_g1 and known house-keeping gene Ribosomal Protein L28 (Rpl28)-Dm01804541_g1.

### *Drosophila* brain dissections, in situ hybridization, and amplification for HCR

For HCR assays, adult heads were fixed in PFA-PBSTx0.1 (1xPBS, 0.1% Triton X-100, 4% PFA) for 20 min while rocking, then washed 3 X 2 min with PBSTx0.01. Brains were dissected in PBSTx0.01 on ice, post-fixed rocking at RT in PFA-PBSTx0.1 for 20 min, and washed rocking at RT in PBSTx0.1 for 3 X 2 min. Brains were further permeabilized in PBSTx0.5 (1X PBS + 0.5% Triton X-100) rocking for 20 min at RT for optimal HCR reagent penetration. HCR v2.0 hybridization and amplification was adapted from previously described protocols (https://files.molecularinstruments.com/MI-Protocol-RNAFISH-FruitFly-Rev7.pdf; Choi et al., 2014; Choi et al., 2018) with the following minor modifications: After permeabilization with PBSTx0.5, brains were pre-hybridized for 20 min at 37°C in preheated hybridization buffer (Molecular Instruments), incubated with NijA-specific probes (Figure S2) diluted to 16 nM in hybridization buffer overnight at 37°C, then washed 4 X 15 min with preheated wash buffer (Molecular Instruments) at 37°C. Brains were washed 2 X 5 min with room temperature 5X SCCT (5x sodium chloride sodium citrate (SSC)/0.1% Tween 20) and pre-amplified in amplification buffer (Molecular Instruments) for 10 min while rocking. HCR Hairpins (Molecular Instruments) were snap-cooled in amplification buffer before being applied. Incubation time for amplification (3 hours at RT) was determined empirically to optimize signal to noise ratio. Brains were washed in 5X SCCT (5 min, 2 X 30 min, and 1 X 5 min) and slide mounted in Vectashield under #1.5 coverslips for confocal imaging.

### Probe design and synthesis

FISH probes against *NijA* were designed using Oligostan (Tsanov et al., 2016). Each gene specific probe was appended at the 5’ end with a full length B2 HCR initiator sequence and a spacer (AAAAA) between the B2 initiator and gene specific sequence (Figure S2). A total of 25 probes were synthesized at 50 pmols each via the oPool platform (Integrated DNA Technologies). Validation of probe specificity was confirmed using *NijAD3* mutants and *in vivo* knockdown of *NijA* by RNAi.

## CRediT authorship contribution statement

**Cole R. Brashaw:** Writing – review and editing, Writing – original draft, Conceptualization, Formal analysis, Investigation, Methodology, Supervision. **Annie M. Griffin:** Writing – review and editing, Formal analysis, Investigation, Methodology. **Sean D. Speese:** Writing – review and editing, Resources, Supervision. **Mary A. Logan:** Writing – review and editing, Writing – original draft, Conceptualization, Funding acquisition, Methodology, Project administration, Supervision.

## Acknowledgements

We would like to thank Andrea Page-McCaw, Dirk Bohmann, and Marc Freeman for generously sharing fly lines. We thank Petra Richer for technical advice on HCR experiments. We thank Tobias Stork and Leire Abalde-Atristain for consulting and technical training. Draper antibody (8A1 RRID:AB_2618106 and 5D14 RRID:AB_2618105) generated by our lab is available at the Developmental Studies Hybridoma Bank, created by the NICHD of the NIH and maintained at The University of Iowa, Department of Biology. Fly stocks obtained from the Bloomington Drosophila Stock Center (NIH P40OD018537) were used in this study. The authors would also like to acknowledge the National Institute of Health (R01NS117934, R21NS107771, R21NS084112) and funds from OHSU Neurology Foundation Funds for generous support of our work.

## Data Statement

Data will be available upon request.

---

**Supplemental Figure 1. Validation of NijA probeset specificity and NijA genetic mutants.**

**(A)** Representative images of ALs from control probe-set only and control hairpin only treatment, 4 hours post antennal nerve injury. Genotype: *w^1118^*. Scale bar = 30µm.

**(B)** RT-qPCR analysis of *NijA* transcript in central brains. 2^−ΔΔCt^ ± SD values are plotted. Mean ± SD plotted; *p=0.0446, ****p<0.0001; ns= not significant; N=4 biological replicates per group. Unpaired t-test for *NijAD3* null mutants. Ordinary one-way ANOVA with Dunnett’s *post-hoc* test for RNAi groups. Genotypes: control = *w^1118^*. NijAD3 = *w^1118^*; *NijAD3/NijAD3*. repo control = *w^1118^; repo-Gal4/+.* NijA RNAi (50632) = *w^1118^*; UAS-NijA^RNAi50632^/+; repo-Gal4/+. NijA RNAi (51358) = *w^1118^; repo-Gal4/ UAS-NijA^RNAi51358^*.

**Supplemental Figure 2. *NijA* B2 HCR probe sequences.**

List of *NijA* transcript-specific sequences (purple) tagged with B2 initiator sequence (red) for HCR amplification.

