## Supplementary material for "Nerve injury promotes glial immune responses through a Draper/Ninjurin A pathway": Supp Figure 1

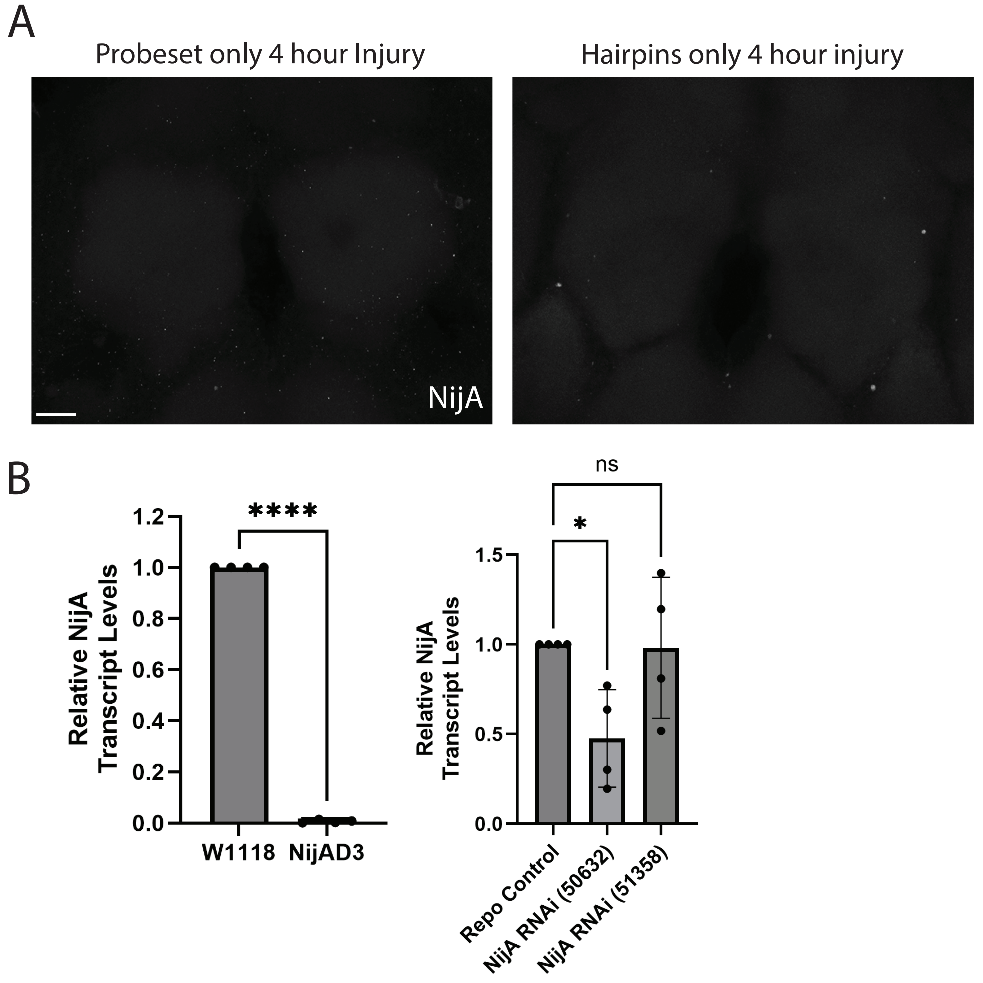


**Supplemental Figure 1. Validation of NijA probeset specificity and NijA genetic mutants.**
(A) Representative images of ALs from control probe-set only and control hairpin only treatment, 4 hours post antennal nerve injury. Genotype: $w^{1118}$. Scale bar = 30μm.
(B) RT-qPCR analysis of *NijA* transcript in central brains. $2^{-\Delta\Delta Ct}$ ± SD values are plotted. Mean ± SD plotted; *p=0.0446, ****p<0.0001; ns= not significant; N=4 biological replicates per group. Unpaired t-test for *NijAD3* null mutants. Ordinary one-way ANOVA with Dunnett's *post-hoc* test for RNAi groups. Genotypes: control = $w^{1118}$. NijAD3 = $w^{1118}$*; NijAD3/NijAD3*. repo control = $w^{1118}$*; repo-Gal4/+*. NijA RNAi (50632) = $w^{1118}$*; UAS-NijA*$^{RNAi50632}$*/+; repo-Gal4/+*. NijA RNAi (51358) = $w^{1118}$*; repo-Gal4/UAS-NijA*$^{RNAi51358}$.
