## Supplementary material for "Nerve injury promotes glial immune responses through a Draper/Ninjurin A pathway": Supp Figure 2

| Probe | Sequence (5' - 3') |
|---|---|
| NijA HCR-B2-1 | CCTCGTAAATCCTCATCAATCATCCAGTAAACCGCCAAAAATTGCCATCTTTAACAAAGGGACGATCGTCGTT |
| NijA HCR-B2-2 | CCTCGTAAATCCTCATCAATCATCCAGTAAACCGCCAAAAATCCAGCGTAATATGCTCTAGGTTACTCATTT |
| NijA HCR-B2-3 | CCTCGTAAATCCTCATCAATCATCCAGTAAACCGCCAAAAAACTGTATAATTATTGATCCTGTTCGCCCGG |
| NijA HCR-B2-4 | CCTCGTAAATCCTCATCAATCATCCAGTAAACCGCCAAAAAATATCGTGACCGTTTTTGATGTTATACTGGCC |
| NijA HCR-B2-5 | CCTCGTAAATCCTCATCAATCATCCAGTAAACCGCCAAAAACATCATTCCCTGAGCCAGGGTCTTCT |
| NijA HCR-B2-6 | CCTCGTAAATCCTCATCAATCATCCAGTAAACCGCCAAAAATGCTGATATGCATTCACATCGGGTATGGG |
| NijA HCR-B2-7 | CCTCGTAAATCCTCATCAATCATCCAGTAAACCGCCAAAAACATCATCATCGGTTTCTGGAACTGCTCT |
| NijA HCR-B2-8 | CCTCGTAAATCCTCATCAATCATCCAGTAAACCGCCAAAAAGCACTCGAAGAGTATTACGTGTCCGTCCG |
| NijA HCR-B2-9 | CCTCGTAAATCCTCATCAATCATCCAGTAAACCGCCAAAAAAGTTTTATTATCTCCTAGTGGCACTTTATCCA |
| NijA HCR-B2-10 | CCTCGTAAATCCTCATCAATCATCCAGTAAACCGCCAAAAAATTTCTTTGTATCTCTTTATCTAGACCTTCTC |
| NijA HCR-B2-11 | CCTCGTAAATCCTCATCAATCATCCAGTAAACCGCCAAAAATTGTTTAAAAACTCGTGCGCAATTTGCGACCG |
| NijA HCR-B2-12 | CCTCGTAAATCCTCATCAATCATCCAGTAAACCGCCAAAAATAAACTATGTTGTATTGGCGGGTAAAGCAGGC |
| NijA HCR-B2-13 | CCTCGTAAATCCTCATCAATCATCCAGTAAACCGCCAAAAAAGTATCTCTGTCCACAGTAAAGGCTGATATTA |
| NijA HCR-B2-14 | CCTCGTAAATCCTCATCAATCATCCAGTAAACCGCCAAAAACAATATAAGGCCCACGCCCACAGCAA |
| NijA HCR-B2-15 | CCTCGTAAATCCTCATCAATCATCCAGTAAACCGCCAAAAAATATAATGCTGAGTGAGATGAACAGCAGACTG |
| NijA HCR-B2-16 | CCTCGTAAATCCTCATCAATCATCCAGTAAACCGCCAAAAAAAGTAAGGATGTTGTGAGCTCGTCTCCAGAAC |
| NijA HCR-B2-17 | CCTCGTAAATCCTCATCAATCATCCAGTAAACCGCCAAAAAAACGCAATTGATTTGCATTCGCGGAGAGAAGT |
| NijA HCR-B2-18 | CCTCGTAAATCCTCATCAATCATCCAGTAAACCGCCAAAAACTCCACGCCGTTGATCGTCACAAAACC |
| NijA HCR-B2-19 | CCTCGTAAATCCTCATCAATCATCCAGTAAACCGCCAAAAAGTCCATTGTATCCCGGAAAGGAGAAACTGGG |
| NijA HCR-B2-20 | CCTCGTAAATCCTCATCAATCATCCAGTAAACCGCCAAAAAGTCTTCCTCCATTGGGAACGTTTACATTCA |
| NijA HCR-B2-21 | CCTCGTAAATCCTCATCAATCATCCAGTAAACCGCCAAAAACGTTGCCTCCGTTGCCTCCAACAGTT |
| NijA HCR-B2-22 | CCTCGTAAATCCTCATCAATCATCCAGTAAACCGCCAAAAAGATTGCACCGCCATAGCTGTGTTTGC |
| NijA HCR-B2-23 | CCTCGTAAATCCTCATCAATCATCCAGTAAACCGCCAAAAATTTGGGCAATTACAAATGAGCAATTCAATGCC |
| NijA HCR-B2-24 | CCTCGTAAATCCTCATCAATCATCCAGTAAACCGCCAAAAAACATTGACCACAGTGACTATAAAAATGCCGC |
| NijA HCR-B2-25 | CCTCGTAAATCCTCATCAATCATCCAGTAAACCGCCAAAAAAAAAAGACCGTCATCCACGCCGGGATTAT |

B2 initiator sequence | spacer | NijA mRNA probe

**Supplemental Figure 2. NijA B2 HCR probe sequences.**
List of *NijA* transcript-specific sequences (purple) tagged with B2 initiator sequence (red) for HCR amplification.
